# Misfolded amyloid β-42 induced impairment of the endosomal-lysosomal pathway revealed by real-time optical monitoring

**DOI:** 10.1101/598540

**Authors:** Karen E Marshall, Devkee Vadukul, Kevin Staras, Louise C Serpell

## Abstract

Misfolding and aggregation of proteins is strongly linked to several neurodegenerative diseases but how such species bring about their cytotoxic actions remains poorly understood. Here we used specifically-designed optical reporter probes and live fluorescence imaging in primary hippocampal neurons to characterise the mechanism by which prefibrillar, oligomeric forms of the Alzheimer’s-associated peptide, Aβ42, exert their detrimental effects. We used a pH-sensitive reporter, Aβ42-CypHer, to track Aβ internalisation in real time, demonstrating that oligomers are rapidly taken up into cells in a dynamin-dependent manner, and trafficked via the endo-lysosomal pathway resulting in accumulation in lysosomes. In contrast, a non-assembling variant of Aβ42 (vAβ42) assayed in the same way is not internalised. Tracking ovalbumin uptake into cells using CypHer or Alexa Fluor tags shows that preincubation with Aβ42 significantly disrupts protein uptake. Our results identify a potential mechanism by which amyloidogenic aggregates directly impair cellular function through disruption of the endosomal-lysosomal pathway.

## INTRODUCTION

Neurodegeneration occurs in several diseases that are characterised by protein aggregation, including Alzheimer’s, Parkinson’s and Huntington’s disease. In each, there is a strong association between the self-assembly of a particular protein to form amyloid fibrils and the development or progression of the disease [1–3]. However, although this ordered self-assembly is tightly linked with cell death, precisely what constitutes a “toxic” structure, and which molecular mechanisms drive detrimental effects, are still unresolved issues.

In Alzheimer’s disease (AD), the pathological state is characterised both by the deposition of tau in neurofibrillary tangles, and Amyloid-β (Aβ) into amyloid plaques. The self-assembly and accumulation of Aβ, in particular the 42-residue form [4, 5], is thought to be a very early event in the progression of Alzheimer’s disease, occurring before changes in tau take place or the expression of cognitive impairment [6, 7]. In support of this, compelling genetic evidence links Aβ aggregation to AD [8], and prefibrillar oligomers of Aβ42 are implicated as critical triggers of neuronal cell death [9, 10, 11]. Additionally, previous studies have shown that synthetic oligomers of Aβ42 cause synaptic dysfunction and neuronal death in a range of model systems both *in vitro* and *in vivo* [1, 12–19].

What is lacking in the current models, however, is a clear understanding of the pathway that links Aβ42 aggregation with the compromised cell health associated with the disease state. Recent work offers an important insight; specifically the observation that proteostasis mechanisms normally operating in low pH lysosomes to degrade macromolecules and maintain cellular health, appear to be defective in AD and other disease states [20–30]. This raises the possibility that Aβ might be specifically exerting its toxicity through its ability to internalise and affect lysosomal pathways.

Here we investigate this key idea directly, using live fluorescence imaging methods both to characterise the uptake and localisation of Aβ42 oligomers in primary hippocampal neurons and to assess their impact on the endosomal-lysosomal pathway. In particular, we exploit a new pH-sensitive optical reporter to show that Aβ oligomers rapidly internalise into cells in a dynamin-dependent manner, and then traffic directly to lysosomes where they remain for several days. Protein misfolding is critical for this outcome since a non-assembling Aβ variant undergoes minimal uptake and accumulation with the same assay. By imaging events occurring in real-time using fluorescent probes, we also demonstrate that the presence of Aβ42 significantly disrupts endocytosis of other proteins. Our findings provide a key link between Aβ42 aggregation and deficits in the endosomal-lysosomal pathway, which may account for early events in disease-related neuronal dysfunction.

## RESULTS

### Toxic Aβ42 oligomers are internalised in defined intracellular compartments

Self-assembly of the Aβ42 peptide into amyloid fibrils is strongly linked with neurodegeneration and the subsequent symptoms observed in AD [1]. As a starting point, we evaluated the time and dose dependent effect of oligomeric Aβ42 [15] on cell viability in primary hippocampal neurons using the Readyprobes assay (Fig. 1a,b). Compared to buffer control, significant differences were observed after incubation with 10 μM Aβ oligomers at all time points, with an increase in cell death from 26.0 ± 3.7% (n total = 1005, dead = 241) to 65.2 ± 8.2% (n total = 568, dead = 369) (Fig. 1b) from 24 hours to 14 days incubation respectively. Previous work using an Aβ control peptide at the same concentration showed that toxic effects are specific to wild type Aβ1-42 [15]. To examine the pathway of action that leads to neuronal cell death, we selected one of the intermediate timepoints (3 d, 44% cell death) for further experiments.

**Figure 1.**
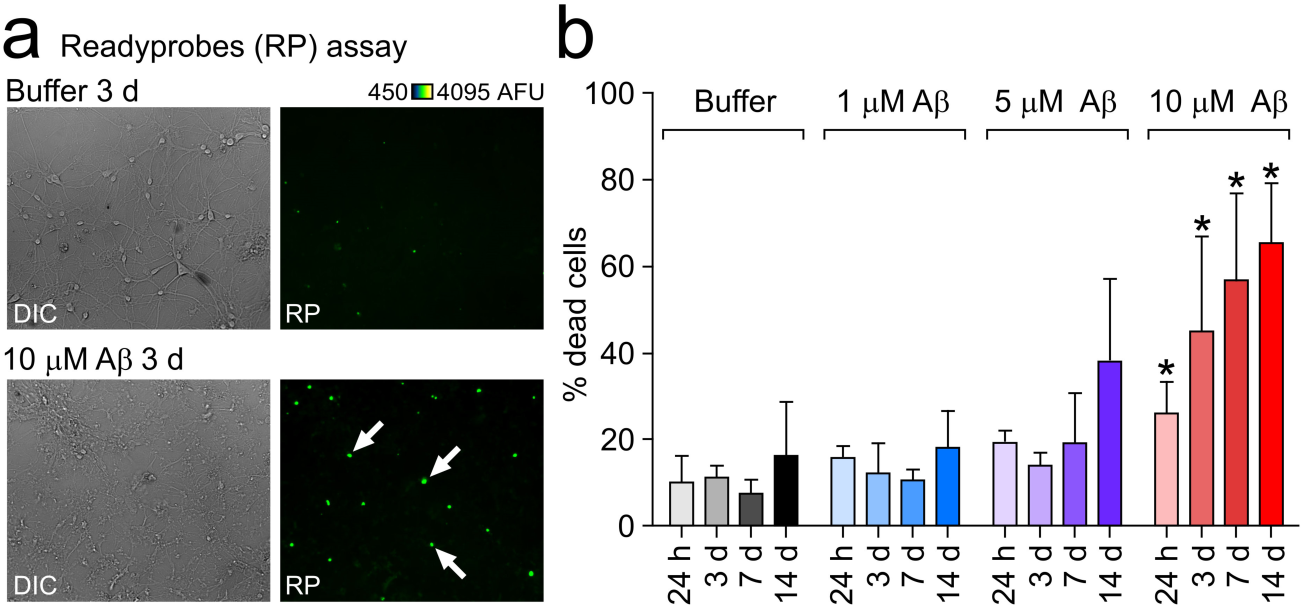
Aβ42 oligomers cause death of primary neurons in a dose and time-dependent manner. (a) Example widefield images following treatment of cells incubated either with buffer or 10 μM Aβ42 oligomers for 3 days with Readyprobes reagent. Left panels are brightfield images showing deteriorated cell morphology in oligomer treated cultures. Right panels are the same region through the green filter, showing dead cells labelled with the green reagent, which stains the nuclei of cells with compromised plasma membrane integrity. More green i.e. dead cells can be seen in oligomer treated cultures (white arrows). (b) Assessment of oligomer toxicity by Readyprobes cell viability assay. Data shown are averages from three FOV (fields of view) in at least three experiments ± SEM. Unpaired student’s t-test; significant differences versus buffer are indicated by *.

Initially, we monitored real-time uptake and trafficking of Alexa Fluor-conjugated Aβ42 oligomers [15, 16]. Importantly, our previous work has demonstrated that Alexa Fluor labelling does not alter the self-assembly of Aβ42 into oligomers or amyloid fibrils [16] and that the fluorescently labelled particles are detected by anti-Aβ antibodies [15]. Fig. 2a and supplementary movie 1 show the uptake of Alexa Fluor 488 labelled Aβ42 oligomers into primary neurons over 15 h. Small fluorescent puncta could be readily observed at the 1 h timepoint (Fig. 2a, arrows) and by 5 hours these were clearly visible in the cytoplasm suggesting that oligomers were being internalised via the plasma membrane into intracellular compartments. This continued in a time-dependent manner (Fig. 2a-c) and at the 3 d incubation timepoint, there was extensive accumulation (Fig. 2b, arrows) with 100% of cells (n = 72) containing visible Aβ42-488 puncta in the cell cytoplasm. Additionally, we saw evidence of fluorescence labelling along the cell boundaries (Fig. 2b, orange arrowheads), indicative of binding of Aβ42-488 oligomers to the extracellular membrane.

**Figure 2.**
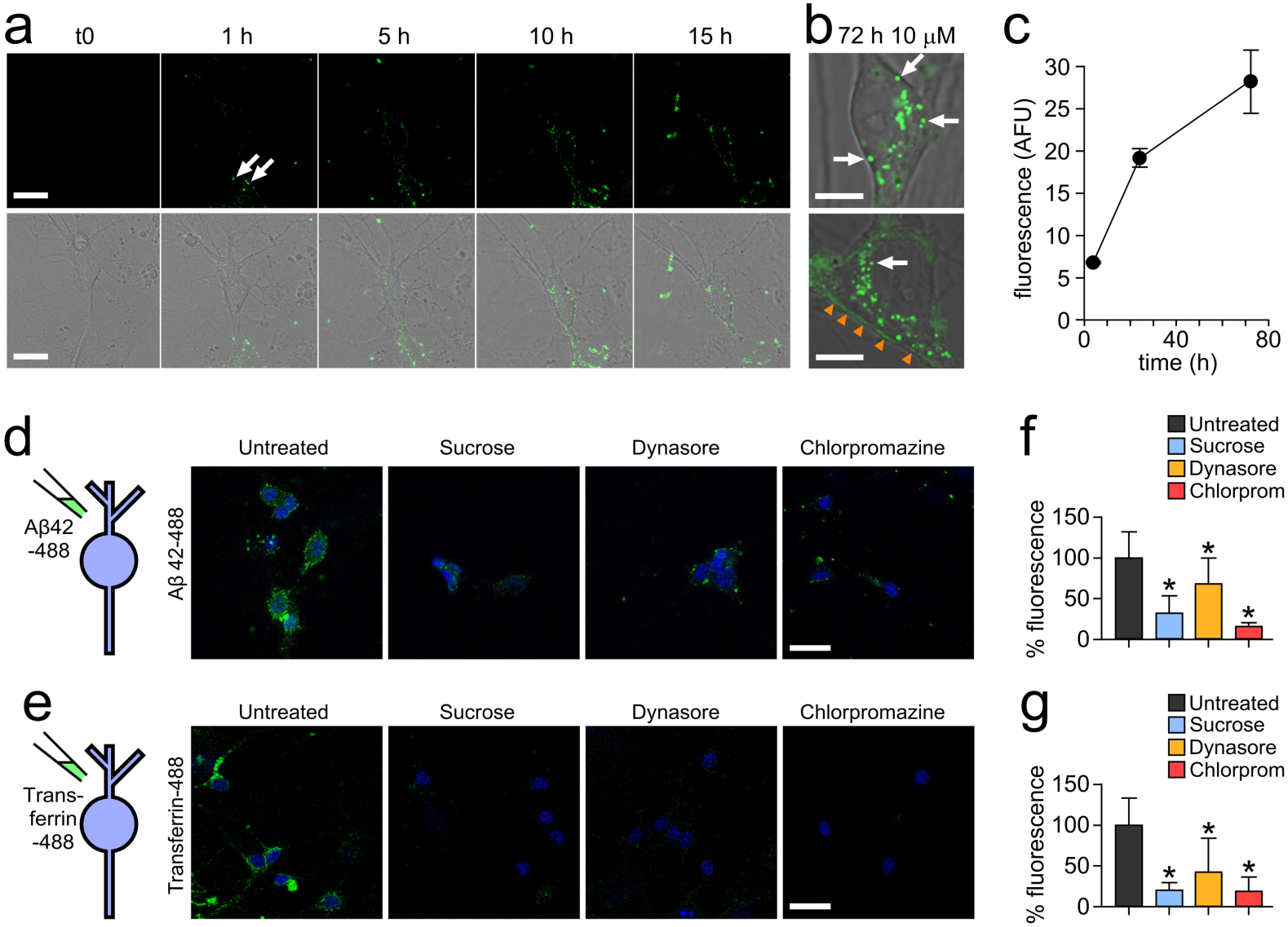
Alexa fluor-488 labelled Aβ42 oligomers are internalised into primary neurons by a dynamin-dependent mechanism. (a) Confocal images showing one 0.5 μm z-slice from the middle of the cell body of neurons treated with 10 μM Aβ42 oligomers over 15 hours. Oligomers can be seen associating with the plasma membrane (white arrows) and internalising into the cell. Scale bar is 25 μm. (b) Magnified example images of cell bodies after 72 hours containing Aβ42-488 oligomers in vesicles within the cytoplasm (white arrows) and along the plasma membrane (orange arrowheads). One 0.5 μm z-slice is shown from the middle of the cell body. Scale bar is 10 μm. (c) Increase in Alexa Fluor 488 fluorescence intensity within the cell body over time. n (cells) 4 hr = 14, 24 hr = 31, 72 hr = 11. Plot shows mean ± SEM. Inhibition of uptake of Aβ42-488 oligomers (d) or transferrin-488 (e) using sucrose, dynasore and chlorpromazine. Neuronal nuclei are shown in blue and Aβ42 or transferrin are green. One 0.5 μm z-slice is shown from the middle of the cell body. Scale bar is 25 μm. (f) and (g) Quantification of 488 fluorescence intensity from the cell body from three experiments normalised to untreated. A Kruskal-Wallis ANOVA was carried out followed by pairwise comparisons using Dunn’s test and all were significant compared to untreated (*).

### Aβ42 internalises in a dynamin-dependent manner into lysosomes

To determine the mechanism of Aβ42-488 oligomer uptake, we tested whether the observed internalisation is sensitive to endocytic blockers using sucrose and chlorpromazine as generic, non-specific inhibitors, and dynasore as a specific inhibitor of the GTP-ase, dynamin. It is well-established that the protein, transferrin, endocytoses via clathrin-coated pits [31, 32] and this was used as a control protein. Serum-starved cells incubated for 15 mins with Aβ42-488 oligomers alone showed bright punctate and diffuse labelling indicative of their association with neurons and internalisation (Fig. 2d, left panel). However, pre-treatment with any of the endocytic inhibitors yielded a striking reduction in fluorescence (Fig. 2d,f) similar to the pattern of distribution in the positive control, transferrin-488 (Fig. 2e,g). These results suggest that Aβ42 oligomers, like transferrin, are endocytosed by a dynamin-dependent mechanism.

Next, we investigated the nature of the cellular structures that accumulate Aβ42 oligomers. To confirm the destination for Aβ42 oligomers following endocytosis, neurons were treated with Alexa Fluor 488-labelled Aβ42 oligomers and co-labelled with Lysotracker red (Fig. 3a), an established marker of lysosomes. As a control, we used a fluorescent conjugate of ovalbumin, a protein known to traffic through the endosomal-lysosomal pathway [33] (Fig. 3b). We found that both protein signals co-localised with the lysotracker fluorescence in compartments around the nucleus, providing clear support for the hypothesis that lysosomes are the final destination for Aβ42 oligomers. As support for this conclusion, we also examined neurons treated with Aβ42 oligomers using transmission electron microscopy (TEM). Aligned with our fluorescence data we observe the presence of large (>1 μm diameter) round electron-dense vesicular structures in the cell cytoplasm (Fig. 3c, arrows). These structures, which were not seen in control neurons, were consistent in appearance with enlarged lysosomal structures reported previously in dystrophic AD neurons [22, 34–36]. Taken together, these abnormalities, specific to oligomer-treated cells, provide further evidence that Aβ42 oligomers have a detrimental effect on the cellular pathways that end with lysosomal degradation.

**Figure 3.**
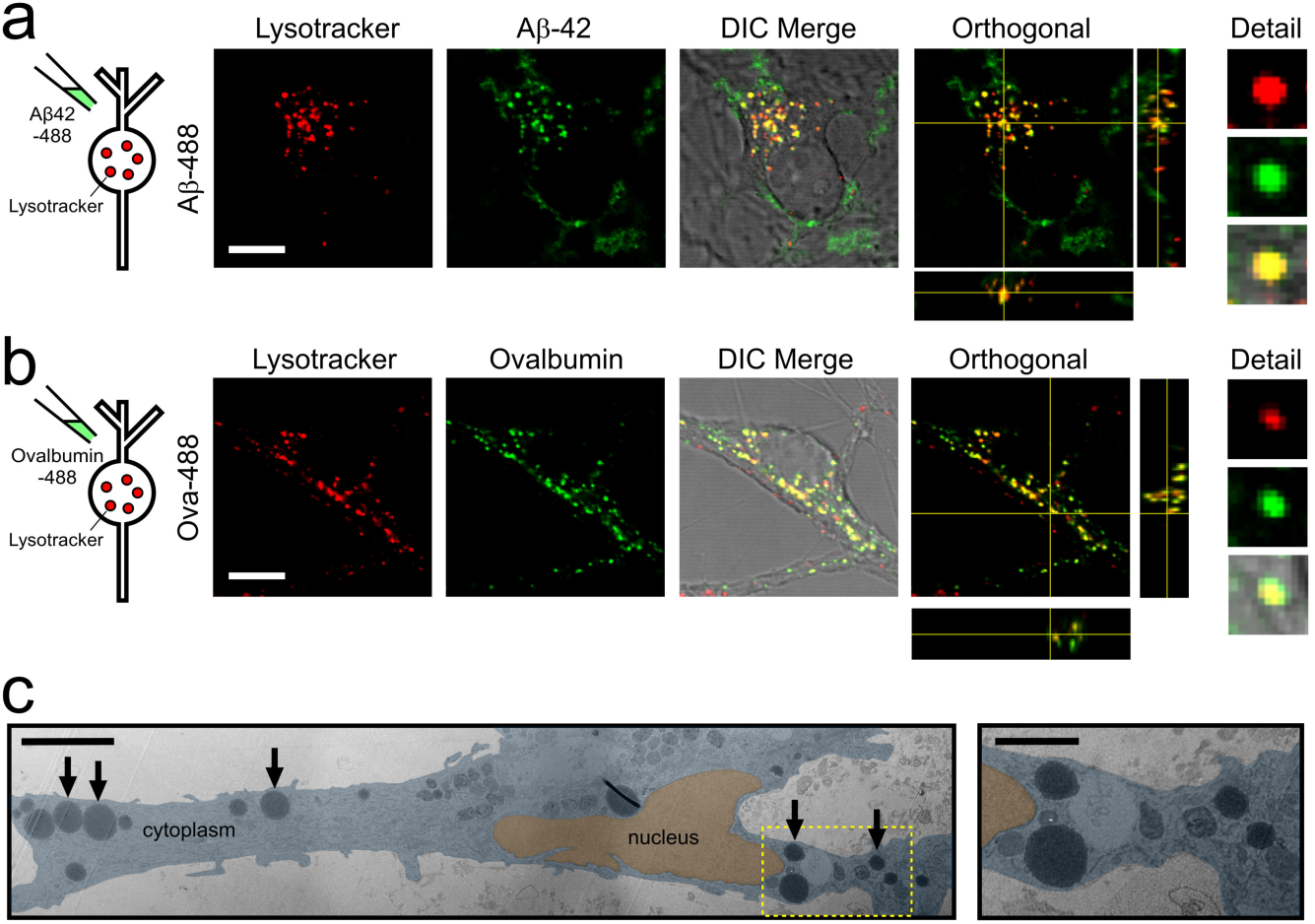
Aβ42 oligomers are trafficked to lysosomes. Co-labelling with Lysotracker red shows that Aβ42-488 oligomers (a) localise to lysosomes with a similar distribution to 488-labelled ovalbumin (b). Orthogonal views also show green and red signal colocalisation in x, y and z. Cells were treated with either 10 μM Aβ42-488 oligomers for 24 hours or 100 μg/mL Ovalbumin-488 for 3 hours. Scale bar is 10 μm. (c) (left panel) Aligned and overlaid transmission electron microscopy images of a section through a neuron treated with 5 μM Aβ42 oligomers for 14 days showing large circular electron-dense structures (arrows). Note, the image is a composite of multiple images that are differentially contrasted to allow clear visualisation of cellular structures. A higher magnification of one region of interest (yellow box) is shown in the right panel. Scale bar, 5 μm (inset 2 μm).

### A novel pH-dependent probe provides a real-time assay of Aβ42 uptake into lysosomes

Next, to provide detailed temporal information on the lysosomal fate of oligomers, we developed a novel optical reporter that provides a real-time readout of Aβ internalisation into acidic compartments. Specifically, we used CypHer5E, a commercially available pH-sensitive fluorescent probe, which exhibits maximal emission when present in an acidic environment [37]. In previous studies this fluorescent moiety has been used to label antibodies for reporting receptor internalisation [38] and to tag vesicle proteins for assaying exo-endocytic recycling at synaptic terminals [39–41].

The endo-lysosomal pathway is characterised by a shift in pH, reducing from ∼6-7 in early endosomes to ∼5 in the lysosome [42, 43]. Therefore, we hypothesised that a protein tagged to CypHer5E that experienced this environmental change would yield important temporal information on internalisation seen as an increase in fluorescence emission, and thus allow us to dynamically monitor uptake.

To validate this approach, we first confirmed that the addition of unconjugated dye to primary hippocampal neurons did not lead to an increase in fluorescence signal (data not shown). As an additional verification, we conjugated CypHer5E with ovalbumin (referred to as Ova-Cy) (Fig. 4a). When Ova-Cy was applied to neurons, we observed a significant time-dependent increase in punctate fluorescence localised to cell bodies, consistent with the presence of this reporter in a cellular compartment that undergoes a drop in pH (Fig. 4b,d). To corroborate this finding, we also tested the effect of pre-treating with NH_4_Cl, a weak base that increases the pH in the lysosome, prior to the addition of Ova-Cy. In this case, a fluorescence increase was not observed, confirming that CypHer5E is an effective monitor of basal lysosomal acidification state (Fig. 4c,d). Moreover, the fluorescence was reduced when NH_4_Cl or bafilomycin, an ATPase blocker, were added to cells following addition of Ova-Cy (data not shown). The pH-dependence of Ova-Cy emission was confirmed *in vitro* using fluorimetry (Fig. 4e). Specifically, this showed that protonated CypHer5E (at low pH) had increased fluorescence at the emission maximum (663 nm) compared with the non-protonated form (at higher pH) when excited with 644 nm light, and vice versa, when excited at 500 nm (the absorbance maximum for CypHer5E at physiological pH). Taken together, our results confirm that CypHer5E is an effective optical reporter for monitoring uptake of target proteins into acidified structures.

**Figure 4.**
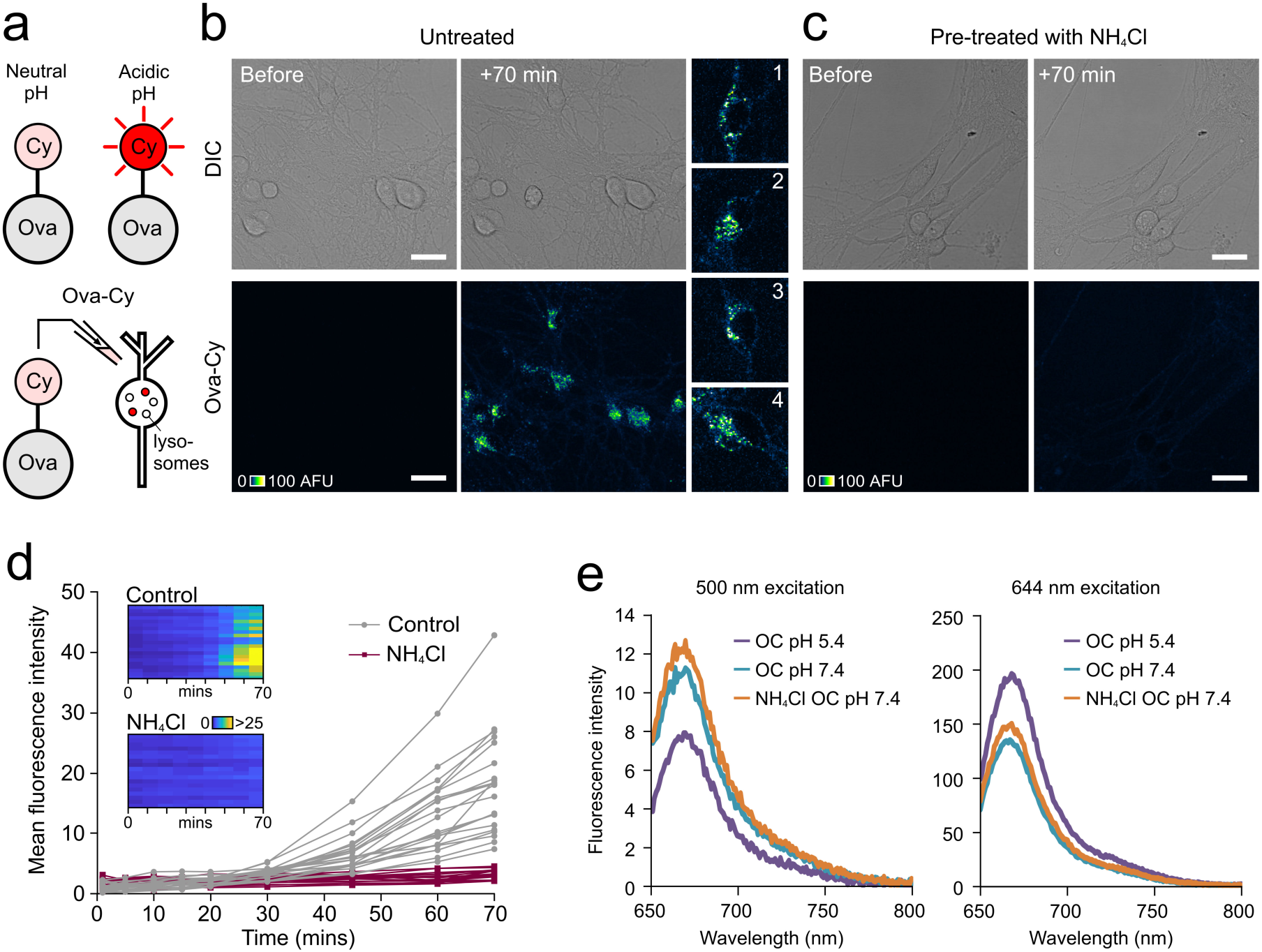
Protein-conjugated CypHer5E reports its pH environment. (a) CypHer5E conjugated to ovalbumin fluoresces upon uptake into intracellular acidic compartments. (b) Confocal images showing increase in fluorescence 70 minutes after addition of Ova-Cy to primary hippocampal neurons. (c) No fluorescence increase is observed in cells pre-treated with NH_4_Cl, a weak base that raises the pH of acidic compartments. One 1.0 μm slice is shown. Scale bar is 25 μm. (d) Quantification of mean fluorescence intensity from cells shown in b and c. Each trace represents an individual cell body either untreated (grey, n = 21) or pre-treated with 10 mM NH_4_Cl for 30 minutes (purple, n = 18). Data pooled from two independent experiments, 3-4 FOV per experiment. (e) Fluorescence arising from Ova-Cy diluted into buffer of pH 5.4 (purple) or 7.4 (turquoise). Ova-Cy diluted into buffer containing NH_4_Cl, pH 7.4 is also shown (orange). The absorbance maximum for CypHer5E at physiological pH 7.4 is 500 nm (left plot) and at pH 5.4 (similar to the pH in a lysosome) is approximately 645 nm. Ova-Cy in acidic (pH 5.4) buffer has increased intensity over neutral pH when excited with 644 nm light. The presence of NH_4_Cl has no effect on the spectral properties of the dye.

Next, we used the same tagging method to examine the fate of internalised Aβ (Fig. 5). Aβ42 was conjugated to CypHer5E (Aβ42-Cy, Fig. 5a and supplementary movie 2) and applied to neurons, resulting in an increase in fluorescence intensity over a 150 min period, comparable to our findings with Ova-Cy (Fig. 5a,c,e). Notably, the distribution of fluorescent puncta in the cytoplasm was consistent with our observations for both Ova-Cy (Fig. 4b) and Aβ42-488 (Figs. 2b and Fig. 3a) suggesting that all were localised in the same way.

**Figure 5.**
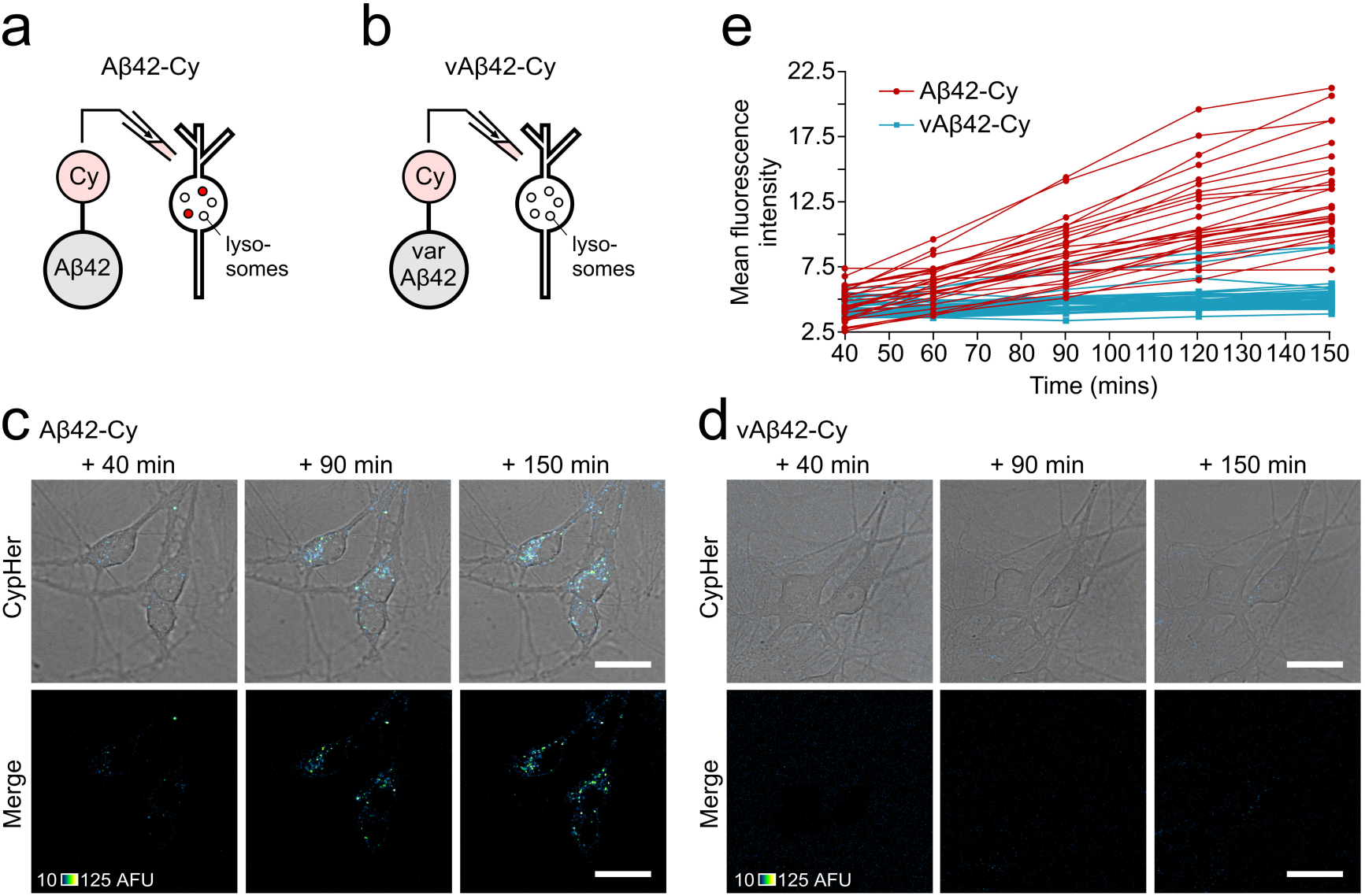
Live tracking of internalisation of self-assembled Aβ42-Cy oligomers into primary neurons and localisation to acidic compartments. a) and b) Cypher5E was conjugated to either Aβ42 or vAβ42 peptides and incubated for 2 hours, during which time Aβ42 is able to self-assemble into oligomers whereas vAβ42 does not. Following addition to primary neurons an increase in fluorescence intensity is observed only in cells treated with 10 μM Aβ42-Cy (c) and not 10 μM vAβ42-Cy (d). One 0.5 μm slice is shown. Scale bar is 25 μm. e) Quantification of mean fluorescence intensity from Aβ42-Cy (red) and Var-Cy (blue) in the cell body (one 0.5 μm slice) over 150 minutes. Each trace arises from the same individual cell. Data pooled from three separate experiments, 4 FOV per experiment, n (cells) Aβ42-Cy = 27, Var-Cy = 42.

To examine whether uptake is specific to prefibrillar misfolded forms of Aβ42, or alternatively a generic outcome with Aβ proteins, we took advantage of a non-assembling form of Aβ42 (variant Aβ42 or vAβ42), which we have recently developed as a control peptide [15]. Importantly, we have previously demonstrated that this variant structure has none of the detrimental effects of Aβ42 with regards to neuronal biology or memory formation [15]. As above, CypHer5E was conjugated to vAβ42 and applied to cells and monitored over 150 minutes. Compared to cells treated with Aβ42-Cy, negligible fluorescence increase was observed in cultures treated with vAβ42-Cy and no change was observed over time (Fig 5d,e).

### Aβ42 oligomers inhibit uptake of ovalbumin in primary neurons

We have shown that Ova-Cy fluorescence effectively reports the pH drop that experienced by the protein trafficking through the endosomal system to lysosomes. This provides us with a powerful assay for monitoring the potential impact of Aβ42 oligomers on endosomal-lysosomal pathway function. To investigate this, cells were pre-treated with Aβ42 oligomers before the addition of Ova-Cy. Strikingly, following Aβ42 exposure, the fluorescence intensity of Ova-Cy was significantly lower than cells pre-treated with buffer alone (Fig. 6a-c). These results provide strong support for a mechanism in which Aβ42 oligomers first traffic to lysosomes and then influence their function.

**Figure 6.**
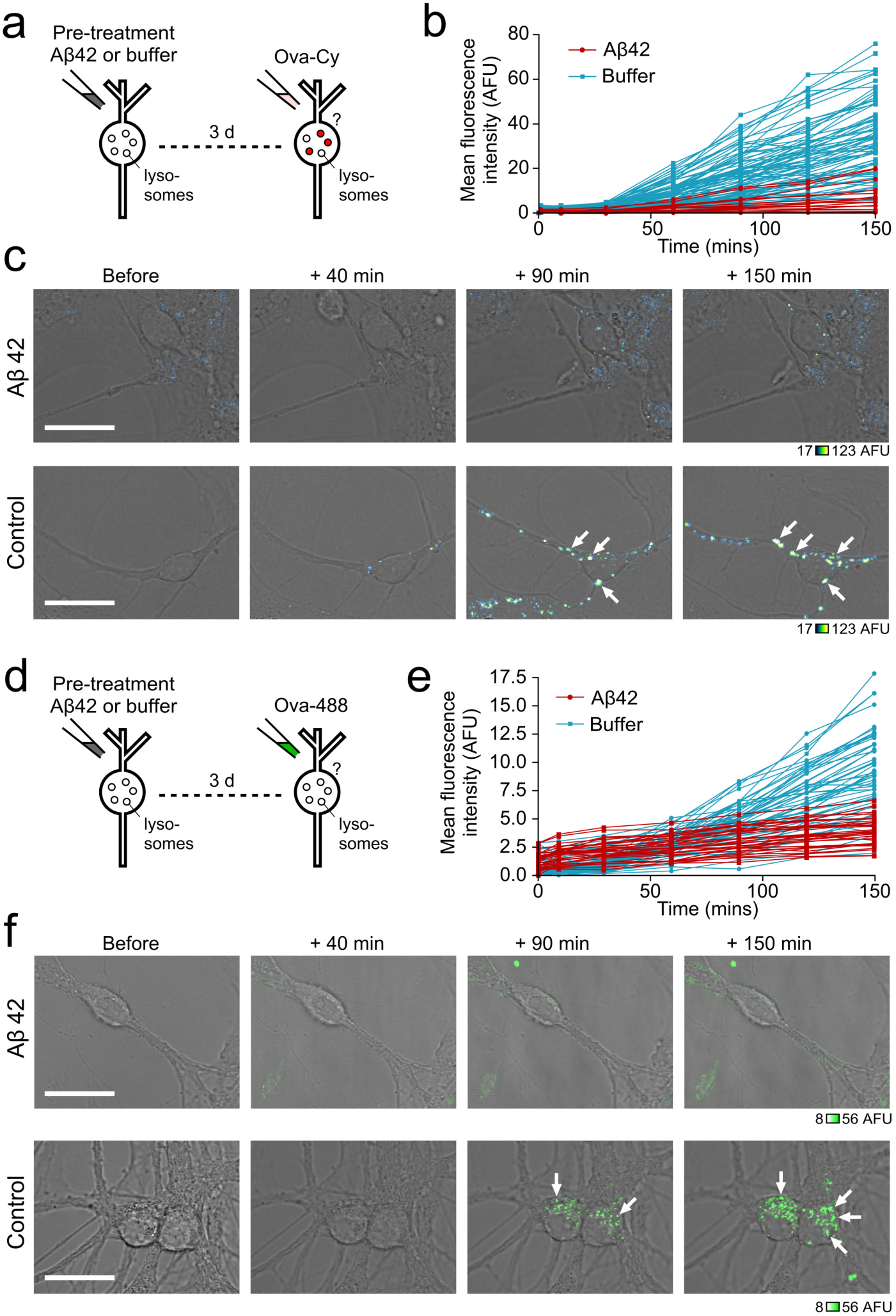
Pre-treating neurons with Aβ42 oligomers inhibits uptake of ovalbumin. Cells were treated either with 10 μM Aβ42 oligomers or equivalent volume of buffer for 72 hours, followed by addition of 25 ug/mL Ova-Cy (a) or Ova-488 (d). Quantification of mean fluorescence intensity from Ova-Cy (b) or Ova-488 (e) in the cell body (one 0.5 μm slice) over 150 minutes from cells either treated with buffer (blue) or pre-treated with 10 μM Aβ42 oligomers for 72 hours (red). Each trace arises from the same individual cell measured over time. Data pooled from three separate experiments 4-9 FOV per experiment, n (cells) buffer = 84, Aβ42 = 19 (b), 4-5 FOV per experiment, n (cells) buffer = 53, Aβ42 = 33 (e). Example images of live cells treated with Aβ42 oligomers (top) or equivalent volume of buffer (bottom) followed by Ova-Cy (c) or Ova-488 (f), taken at increasing time points. Fluorescence intensity is only reduced in cells treated with oligomers. Arrows point to neuronal cell bodies. One 0.5 μm slice is shown. Scale bar is 25 μm.

To support this conclusion, we carried out a complementary experiment in which we tagged ovalbumin with Alexa Fluor 488 (Fig. 6d-f). Analogous to the Ova-Cy experiments, we found a significantly reduced fluorescence intensity in cells pre-treated with Aβ42 oligomers compared to control, showing that the uptake and trafficking of Ova is impaired. Taken together, our findings suggest that the presence of intracellular Aβ42 causes a generalised impairment of the endosomal-lysosomal system likely contributing to the toxic action of Aβ42 on living cell function.

## DISCUSSION

The ordered assembly of peptides and proteins into amyloid is strongly linked with neurodegeneration *in vivo* and *in vitro* [44, 45]. Understanding the cause-and-effect relationship between protein self-assembly and neuronal dysfunction is challenging since early events are difficult to capture or recapitulate. Nevertheless, the case for protein misfolding as a cause of neuronal loss is compelling [11, 46–48]. On the strongly-evidenced premise that misfolded proteins are able to damage neurons, we developed an experimental model using oligomers of the Alzheimer’s-associated Aβ42 peptide in conjunction with novel applications for fluorescent probes, to explore the link between oligomer exposure and degeneration of hippocampal neurons. Benefiting from time-lapse imaging of specially engineered optical reporters in living neurons, we characterised structural and functional properties of cells in real-time, as a powerful strategy to elucidate the underlying mechanisms of aggregated peptide-induced neuronal cell death. Initially, we confirmed that the synthetic, pre-fibrillar Aβ42 oligomers were toxic to primary hippocampal neurons, consistent with previous findings [1, 2, 49]. Using fluorescence imaging, we then examined the endosomal-lysosomal physiology of individual live cells that contained endocytosed Aβ42 oligomers. This real-time readout allowed us to reveal dysfunctional mechanisms prior to cell death, providing important insights into early events that can lead to neurodegeneration. Specifically, we found that oligomers are internalised into neurons predominantly by a dynamin-dependent mechanism and that these are then trafficked to lysosomes and remain intracellular for several days.

It is well-established that the acidification of lysosomes is critical to their function [50]. In order to understand the potentially direct effects of Aβ42 oligomers on lysosomal function, we developed a method to tag Aβ42 and ovalbumin with a pH-sensitive probe (CypHer, Roche) that selectively fluoresces in acidic compartments. Using this approach, the uptake and localisation of Aβ42-Cy peptides within the cell could be monitored in real time and compared with an assembly-incompetent variant of Aβ42. In particular, our findings showed that this uptake is specific to self-assembling, wild-type Aβ42.

To explore the downstream effects of accumulated Aβ42 on the endo-lysosomal pathway, we utilised tagged versions of ovalbumin to assay uptake and acidification events in cells pre-treated with Aβ42 oligomers. We found that the levels of Ova-488 and Ova-Cy fluorescence were significantly reduced in Aβ42 treated cells compared to controls, implying that the accumulated Aβ oligomers are affecting the endo-lysosomal pathway. Because we only considered live cells in this analysis (those with an intact and smooth plasma membrane and a clearly defined nucleus, where often nucleoli were visible), we can be confident in concluding that the impairment of the endosomal-lysosomal system occurs prior to cell death. This implicates a potentially catastrophic and ultimately fatal effect of Aβ42 on neuronal cells.

How does internalised Aβ42 impair the endosomal-lysosomal pathway? We propose that a number of mechanisms could conceivably be involved. These include a direct effect of Aβ42 on acidification through action on the ATPase [26, 27], through lysosomal leakage associated with membrane interactions [30, 51, 52], or simply through its accumulation and overloading of lysosomal proteolytic capacity [53–55]. In our study, we clearly demonstrate that a consequence of Aβ42 internalisation is an inhibition of further endocytosis, implying that the oligomers stall a process critical for cellular health. It is interesting to note the presence of Aβ42 on the extracellular side of the cell membrane, which could provide a site for oligomeric binding, or further aggregation [56]. In either case, it is conceivable that the accumulation of aggregated proteins along the plasma membrane could perturb endocytosis and might be another potential mechanism for our observations.

Collectively these results outline a mechanism of Aβ42-induced cell death that leads to the disrupted endocytosis of essential proteins and nutrients. Indeed, they offer a mechanistic basis for previous *in vivo* studies that have reported defects in lysosomal function in neurodegenerative diseases, including the appearance of abnormal lysosomes and autophagosomes within dystrophic neurites in human AD and mouse model brains [21, 26, 34, 35, 57]. Abnormalities to the endocytic pathways in neurons from AD brain tissue have also been observed [58]. Here we have gained key information on the specific pathway of action by which misfolded proteins exert toxic action on cells whereby uptake of Aβ42 oligomers into neurons renders them unable to carry out further endocytosis. We suggest that the accumulation of aggregated Aβ42 within lysosomes stalls the endosomal system and that this is an early event in oligomer-induced neurodegeneration, and collectively provides an explanation for cytotoxicity caused by misfolded proteins.

## METHODS

### Peptide preparation

#### Aβ42 oligomers and vAβ42

Hexafluoroisopropanol (HFIP) films of recombinant Aβ1-42 were purchased from rPeptide and vAβ1-42 was purchased from JPT Peptide Technologies. Peptides were prepared using an adapted protocol [59], which ensures the removal of potentially contaminating trace solvents. HEPES buffer (10 mM HEPES, 50 mM NaCl, 1.6 mM KCl, 2 mM MgCl_2_, 3.5 mM CaCl_2_) was used to mimic the culture media as previously described [16, 59, 60]. All preparation was conducted using protein LoBind Eppendorfs and tips (Alpha Laboratories). Briefly, 0.2 mg Aβ1-42 (rPeptide) was solubilised in 200 μL HFIP (Sigma-Aldrich) to disaggregate the peptide. The solution was then vortexed on high for one minute and sonicated in a 50/60 Hz bath sonicator for five minutes. HFIP was dried completely using a low stream of nitrogen gas. Once completely dried, 200 μL dry dimethylsulfoxide (DMSO) (Sigma-Aldrich) was added, vortexed for one minute, and sonicated for one minute. Solutions were added to a 7K MWCO Zeba buffer-exchange column (Thermo Scientific) equilibrated with HEPES buffer with 40 μL HEPES as a stacking buffer. The protein solution was kept on ice and the absorbance at 280 nm measured with a NanoDrop spectrophotometer using a molar absorption coefficient of 1490 M^−1^ cm^−1^. Solutions were immediately diluted to 50 μM with HEPES buffer and incubated where indicated or used immediately in further experiments.

#### Alexa Fluor conjugated peptides

Peptides were treated as outlined above up to the addition of 200 μL dry DMSO. Alexa Fluor tags were prepared as per the manufacturer’s instructions (Life Technologies). Briefly, 10 μL dH_2_O was added to the Alexa Fluor TFP ester and kept on ice. This was added to the peptide in 200 μL DMSO along with 20 μL 1M sodium bicarbonate pH 8.3 to increase the reaction efficiency, mixed and incubated for 15 minutes at room temperature protected from light. Following this, the procedure resumed as above, and the solution was then added to the Zeba buffer exchange column. The calculations for the concentration were adjusted to take into account the absorbance of the dye.

### Cell culture

Rats were housed within a specialised facility under Home Office UK guidelines and sacrificed using procedures in accordance with Animals (Scientific Procedures) Act 1986, Amendment Regulations 2012 and approved by a local ethical review panel. Primary neurons were prepared from the hippocampus of P0-P1 rats as described previously [61] by initially dissecting the tissue into ice cold HBSS containing 0.1 M HEPES. Following washes in pre-warmed basal Eagle’s Minimum (BME) (Gibco) containing 0.5% glucose, 2% FCS, 1 mM Na-Pyruvate, 0.01 M HEPES pH 7.35, 1% Penicillin-Streptomycin, 1% B27 supplement and 1% Glutamax, tissues were triturated using a 1 mL pipette until fully dissociated. The cell suspension was diluted further with complete BME media and approximately 40,000 cells plated into 2 cm^2^ wells containing a coverslip coated in 20 μg/mL Poly-D-Lysine with a layer of hippocampal astrocytes that had been growing for 4-5 days. After 2-3 days, cells were treated with 3.25 μM cytosine arabinoside to curb further proliferation of astrocytes. Experiments were carried out on cells 10-14 DIV.

### Cell viability assay

After the desired incubation time, one drop of each ReadyProbes reagent (Life Technologies) was added to cells in 500 μL of media. The kit contains a blue stain to label all cells, and a green stain to label dead cells only. Cells were incubated at 37 °C for 15 minutes and then imaged using widefield fluorescence microscopy with a Zeiss Cell Observer widefield microscope fitted with a Hamamatsu ORCA camera and a Nikon Plan Fluor 10X NA 0.3 air objective. DAPI fluorescence was captured using a G 365 excitation filter and a LP 420 emission filter with a FT 395 dichroic. Green fluorescence was captured using a FITC filter set (BP 450-490 excitation filter, BP 515-565 emission filter and FT 510 dichroic). Identical acquisition settings were used for all experimental and control samples. Images were analysed using FIJI. Neuronal cell bodies were identified using DIC and DAPI channels (astrocytes were excluded), which indicated total cell number (live and dead). Those that were also fluorescent in the FITC channel were defined as dead. The numbers of dead cells were expressed as a percentage of the total number of cells. At least three fields of view were imaged per sample in each experiment. Each experiment i.e. using newly prepared peptide added to different cultures, was performed at least three times.

### Aβ42 oligomer uptake and blocking

All imaging experiments outlined below were carried out using a Leica SP8 confocal microscope and image processing using FIJI. For overnight imaging, the Adaptive Focus Control feature was used to maintain constant focal planes throughout the course of the experiment. Where multiple fluorescent probes were imaged, samples were scanned sequentially to prevent spectral bleedthrough.

To monitor live uptake, neurons were plated on top of a layer of astrocytes in poly-D-Lysine coated 35 mm dishes with 1.5 coverslips (Mattek) and cultured for 10-14 days. Fluorescent-tagged oligomers were added and cells imaged live using a 63x numerical aperture (NA) 1.2 oil objective. The environment was maintained at 37°C with humidified CO_2_.

For blocking experiments, cells plated on coverslips in 24 well plates were washed with warm external bath solution (EBS; 137 mM NaCl, 5 mM KCl, 3 mM CaCl_2_, 1 mM MgCl_2_, 10 mM D-Glucose and 5 mM HEPES pH 7.3) then serum starved for 1 hour in 500 μL warm EBS. To block endocytosis, cells were treated with one of the following for 30 minutes: 0.5 M sucrose, 10 μM Dynasore (Sigma), 25 μg/mL chlorpromazine or left untreated. After blocking, fluorescently labelled Aβ42 oligomers were added, or 100 μg/mL transferrin-488 as a control to confirm blocking effects. After incubation for 30 minutes, cells were washed 3-4 times quickly with warm EBS, fixed using 2% paraformaldehyde (PFA) in EBS for 15 minutes, rinsed with PBS then the coverslip mounted onto a glass slide using Prolong Gold with DAPI (Life Technologies). Fixed cells were imaged with a 40x NA 1.1 water objective.

### Lysosome labelling

Ovalbumin-488 was purchased from Life Technologies and reconstituted in PBS to a stock concentration of 5 mg/mL Hippocampal neurons plated on Mattek dishes were treated with Ova-488 to a final concentration of 100 μg/mL for 3 hours or Aβ42-488 oligomers to 10 μM for 24 hours, then 100 nM Lysotracker red (Life Technologies) was added, incubated for 1.5 hours and imaged using a 100x oil objective.

### Transmission electron microscopy

Cells grown on coverslips were washed once with external bath solution (EBS, 37 mM NaCl, 5 mM KCl, 2.5 mM CaCl_2_, 1 mM MgCl_2_, 10 mM D-glucose, 5 mM HEPES) then fixed in 2% paraformaldehyde, 2% glutaraldehyde in PBS for 15 mins at room temperature. The fixative was replaced with 100 mM glycine and incubated for one hour followed by a 1 min wash in 100 mM NH_4_Cl. Cells were then washed twice with 0.1 M sodium cacodylate buffer before a second fixation step in 1% osmium tetroxide with 1.5 % potassium ferrocyanide in sodium cacodylate for 1 hour. Following repeat rinses with 0.1 M sodium cacodylate, 10 mg/ml tannic acid was added (45 mins) to stabilize and enhance contrast, before a further incubation in 10 mg/ml anhydrous sodium sulphate (5 mins). The cells were rinsed with water (5 repeats) and dehydrated using increasing concentrations of ethanol (50 %, twice with 70 %, twice with 90 % then three times with 100%, 5 minutes each). The coverslips were then transferred into a 1:1 mixture of polypropylene oxide and EPON resin for 1 hour, then moved into pure EPON resin (Electron Microscopy Sciences) (exchanged after 2 hrs) and left overnight. The following day, the coverslip was placed cell-side down onto an EPON block and dried at 60 °C overnight until polymerised. The coverslip was removed by submerging in liquid nitrogen and the block sectioned (70 nm thickness). Sections were viewed on a JEOL transmission electron microscope (100 kV).

### Conjugation of CypHer5E to Ovalbumin and Aβ peptides

CypHer5E Mono NHS Ester (Roche) was prepared as per the manufacturer’s instructions. Briefly, 100 μL of dimethyl sulphoxide (DMSO) was added to 1 mg CypHer5E and the solution sonicated in a 50/60 Hz bath sonicator for 15 seconds. 5 μL of dye was diluted into 4 mL PBS-sodium bicarbonate (0.5 M) pH 8.3 and the absorbance at 500 nm measured. The concentration of dye was calculated based on a molar extinction coefficient of 40,000. Chicken egg white albumin was purchased from Sigma and dissolved to a concentration of 10 mg/mL in water. The concentration was confirmed using absorbance readings at 280 nm and an extinction coefficient of 32,176. The solution was diluted to 1 mg/mL in PBS-sodium bicarbonate (0.5 M) pH 8.3 and a 20 M excess of CypHer5E dye added. The solution was protected from light and gently agitated on a rocker for 1 hour. Following the labelling reaction, unconjugated dye was separated and removed from the solution using a purification column packed with BioGel P-30 fine size exclusion purification resin (BioRad), which is designed to separate free dye from proteins with a molecular weight of greater than 40,000 kDa (MW ovalbumin = 45,000 kDa). Elution buffer (10 mM potassium phosphate, 150 M sodium chloride pH 7.2) was run through the column and once excess buffer had drained into the column bed the reaction mixture was added, allowed to enter into the resin, then eluted using elution buffer. The labelled protein (Ova-Cy) was collected and the concentration of protein and dye measured using a Nanodrop spectrophotometer. The stock solution was kept at 4 °C for a maximum of 3 months. For Aβ42 and vAβ42, labelling with CypHer5E was performed using a protocol similar to the Alexa Fluor labelling method described previously. CypHer5E dye was prepared as described above and a 20 M excess added to the peptide dissolved in 200 μL DMSO along with 20 μL 1 M sodium bicarbonate pH 8.3. Briefly, the reaction mixture was incubated for 15 minutes then applied to a Zeba buffer-exchange column equilibrated with HEPES to remove the DMSO and free dye. The protein and dye concentrations of the eluted solutions were measured, diluted to 50 μM (protein), incubated for 2 hours to form oligomers (for WT Aβ42) and then diluted to working concentration.

### Live-cell imaging of neurons treated with CypHer5E conjugated proteins

Neurons were plated on 35 mm Mattek dishes as described previously. After 10-14 DIV, cells were either treated with Aβ42 oligomers, Ova-Cy, Aβ42-Cy, vAβ42-Cy or an equivalent volume of HEPES buffer. All imaging was carried out live in media at 37°C with humidified CO_2_ using a 1.4 NA HC PL APO CS2 63x oil objective with a 633 nm laser line and fluorescence emission collected between 650 and 700 nm with a PMT detector. For experiments assessing pH dependence of CypHer, Ova-Cy was added to a final concentration of 10 μg/mL with or without 50 mM NH_4_Cl and imaging started. For experiments monitoring a decrease in fluorescence, NH_4_Cl was added to a final concentration of 50 mM or bafilomycin to 1 μM and imaging continued. For experiments where cells were pre-incubated with Aβ42 or vAβ42, Ova-Cy was added to a final concentration of 25 μg/mL. For all CypHer analyses, where one 0.5 μM z-slice has been analysed, a focal plane in the middle of the cell that contained none or a negligible proportion of saturating pixels within the cell body was selected from the image that had maximal fluorescence. Focal planes in images at other time points were matched to this slice to ensure the same focal plane of the cell was being monitored and mean fluorescence intensity calculated. A region of interest was drawn around the cell body excluding the nucleus. The mean fluorescence intensity in either one z-slice or a maximum projection image of several slices at each time point was measured in each cell body. All cells within a given FOV were analysed. Where images are displayed, LUTs were matched for each image within an experiment. All analysis was carried out using FIJI. Graphs and statistical analysis were prepared in GraphPad Prism.

### Fluorimetry

19 mM Ova-Cy stock solution was diluted ten-fold into EBS that had a pH of either 5.4 or 7.4 when it contained 10 mM NH4Cl. Scans were collected on a Varian Cary Eclipse fluorimeter. Samples were excited at either 500 nm or 644 nm and emission collected from 650 to 800 nm. Excitation and emission slit widths were set to 5 nm, and the scan rate was 600 nm/min with 1 nm data intervals and an averaging time of 0.1 seconds.

## Data availability

The datasets supporting the conclusions of this article are included within the article and its supporting information.

## Acknowledgements

LCS, KS and KM are supported by funding from Medical Research Council UK (MR/K022105/1). LCS is supported by Alzheimer’s society and Alzheimer’s research UK. KS is supported by funding from the BBSRC (BB/K019015/1).

## Author contribution statement

KM and DV conducted the toxicity and uptake experiments. KM conducted remaining experiments, managed the work and wrote the manuscript. KM, LS and KS planned the experiments. LS and KS directed the work and edited the manuscript and figures.

## Materials and correspondence

Request for materials may be addressed to LCS or KS.

## References

1. Walsh, D.M., et al., Naturally secreted oligomers of amyloid beta protein potently inhibit hippocampal long-term potentiation in vivo. Nature, 2002. 416(6880): p. 535–9.

2. Kayed, R. and C.A. Lasagna-Reeves, Molecular mechanisms of amyloid oligomers toxicity. J Alzheimers Dis, 2013. 33 Suppl 1: p. S67–78.

3. Glabe, C.G., Common mechanisms of amyloid oligomer pathogenesis in degenerative disease. Neurobiol Aging, 2006. 27(4): p. 570–5.

4. Jarrett, J.T., E.P. Berger, and P.T. Lansbury, Jr., The carboxy terminus of the beta amyloid protein is critical for the seeding of amyloid formation: implications for the pathogenesis of Alzheimer’s disease. Biochemistry, 1993. 32(18): p. 4693–7.

5. Bitan, G., et al., Amyloid beta-protein (Abeta) assembly: Abeta 40 and Abeta 42 oligomerize through distinct pathways. Proc Natl Acad Sci U S A, 2003. 100(1): p. 330–5.

6. Roher, A.E., et al., beta-Amyloid-(1-42) is a major component of cerebrovascular amyloid deposits: implications for the pathology of Alzheimer disease. Proc Natl Acad Sci U S A, 1993. 90(22): p. 10836–40.

7. Jack, C.R., Jr., et al., Tracking pathophysiological processes in Alzheimer’s disease: an updated hypothetical model of dynamic biomarkers. Lancet Neurol, 2013. 12(2): p. 207–16.

8. Scheuner, D., et al., Secreted amyloid beta-protein similar to that in the senile plaques of Alzheimer’s disease is increased in vivo by the presenilin 1 and 2 and APP mutations linked to familial Alzheimer’s disease. Nat Med, 1996. 2(8): p. 864–70.

9. Mc Donald, J.M., et al., The presence of sodium dodecyl sulphate-stable Abeta dimers is strongly associated with Alzheimer-type dementia. Brain, 2010. 133(Pt 5): p. 1328–41.

10. Lesne, S.E., et al., Brain amyloid-beta oligomers in ageing and Alzheimer’s disease. Brain, 2013. 136(Pt 5): p. 1383–98.

11. Hardy, J. and D.J. Selkoe, The amyloid hypothesis of Alzheimer’s disease: progress and problems on the road to therapeutics. Science, 2002. 297(5580): p. 353–6.

12. Snyder, E.M., et al., Regulation of NMDA receptor trafficking by amyloid-beta. Nat Neurosci, 2005. 8(8): p. 1051–8.

13. Lambert, M.P., et al., Diffusible, nonfibrillar ligands derived from Abeta1-42 are potent central nervous system neurotoxins. Proc Natl Acad Sci U S A, 1998. 95(11): p. 6448–53.

14. Cheng, I.H., et al., Accelerating amyloid-beta fibrillization reduces oligomer levels and functional deficits in Alzheimer disease mouse models. J Biol Chem, 2007. 282(33): p. 23818–28.

15. Marshall, K.E., et al., A critical role for the self-assembly of Amyloid-beta1-42 in neurodegeneration. Sci Rep, 2016. 6: p. 30182.

16. Soura, V., et al., Visualization of co-localization in Abeta42-administered neuroblastoma cells reveals lysosome damage and autophagosome accumulation related to cell death. Biochem J, 2012. 441(2): p. 579–90.

17. Lacor, P.N., et al., Synaptic targeting by Alzheimer’s-related amyloid beta oligomers. J Neurosci, 2004. 24(45): p. 10191–200.

18. Shankar, G.M., et al., Natural oligomers of the Alzheimer amyloid-beta protein induce reversible synapse loss by modulating an NMDA-type glutamate receptor-dependent signaling pathway. J Neurosci, 2007. 27(11): p. 2866–75.

19. Lesne, S., et al., A specific amyloid-beta protein assembly in the brain impairs memory. Nature, 2006. 440(7082): p. 352–7.

20. Nixon, R.A., Autophagy, amyloidogenesis and Alzheimer disease. J Cell Sci, 2007. 120(Pt 23): p. 4081–91.

21. Terry, R.D., N.K. Gonatas, and M. Weiss, Ultrastructural Studies in Alzheimer’s Presenile Dementia. Am J Pathol, 1964. 44: p. 269–97.

22. Suzuki, K. and R.D. Terry, Fine structural localization of acid phosphatase in senile plaques in Alzheimer’s presenile dementia. Acta Neuropathol, 1967. 8(3): p. 276–84.

23. Cataldo, A.M., et al., Gene expression and cellular content of cathepsin D in Alzheimer’s disease brain: evidence for early up-regulation of the endosomal-lysosomal system. Neuron, 1995. 14(3): p. 671–80.

24. Nixon, R.A. and A.M. Cataldo, Lysosomal system pathways: genes to neurodegeneration in Alzheimer’s disease. J Alzheimers Dis, 2006. 9(3 Suppl): p. 277–89.

25. Yang, D.S., et al., Reversal of autophagy dysfunction in the TgCRND8 mouse model of Alzheimer’s disease ameliorates amyloid pathologies and memory deficits. Brain, 2011. 134(Pt 1): p. 258–77.

26. Wolfe, D.M., et al., Autophagy failure in Alzheimer’s disease and the role of defective lysosomal acidification. Eur J Neurosci, 2013. 37(12): p. 1949–61.

27. Lee, J.H., et al., Presenilin 1 Maintains Lysosomal Ca(2+) Homeostasis via TRPML1 by Regulating vATPase-Mediated Lysosome Acidification. Cell Rep, 2015. 12(9): p. 1430–44.

28. Gowrishankar, S., Y. Wu, and S.M. Ferguson, Impaired JIP3-dependent axonal lysosome transport promotes amyloid plaque pathology. J Cell Biol, 2017. 216(10): p. 3291–3305.

29. Yang, A.J., et al., Loss of endosomal/lysosomal membrane impermeability is an early event in amyloid Abeta1-42 pathogenesis. J Neurosci Res, 1998. 52(6): p. 691–8.

30. Liu, R.Q., et al., Membrane localization of beta-amyloid 1-42 in lysosomes: a possible mechanism for lysosome labilization. J Biol Chem, 2010. 285(26): p. 19986–96.

31. Harding, C., J. Heuser, and P. Stahl, Receptor-mediated endocytosis of transferrin and recycling of the transferrin receptor in rat reticulocytes. J Cell Biol, 1983. 97(2): p. 329–39.

32. Mayle, K.M., A.M. Le, and D.T. Kamei, The intracellular trafficking pathway of transferrin. Biochim Biophys Acta, 2012. 1820(3): p. 264–81.

33. Zhang, T., et al., Lysosomal cathepsin B plays an important role in antigen processing, while cathepsin D is involved in degradation of the invariant chain inovalbumin-immunized mice. Immunology, 2000. 100(1): p. 13–20.

34. Nixon, R.A., et al., Extensive involvement of autophagy in Alzheimer disease: an immuno-electron microscopy study. J Neuropathol Exp Neurol, 2005. 64(2): p. 113–22.

35. Gowrishankar, S., et al., Massive accumulation of luminal protease-deficient axonal lysosomes at Alzheimer’s disease amyloid plaques. Proc Natl Acad Sci U S A, 2015. 112(28): p. E3699–708.

36. Fazzari, P., et al., PLD3 gene and processing of APP. Nature, 2017. 541(7638): p. E1–E2.

37. Adie, E.J., et al., A pH-sensitive fluor, CypHer 5, used to monitor agonist-induced G protein-coupled receptor internalization in live cells. Biotechniques, 2002. 33(5): p. 1152–4, 1156–7.

38. Riedl, T., et al., High-Throughput Screening for Internalizing Antibodies by Homogeneous Fluorescence Imaging of a pH-Activated Probe. J Biomol Screen, 2016. 21(1): p. 12–23.

39. Kahms, M. and J. Klingauf, Novel pH-Sensitive Lipid Based Exo-Endocytosis Tracers Reveal Fast Intermixing of Synaptic Vesicle Pools. Front Cell Neurosci, 2018. 12: p. 18.

40. Truckenbrodt, S., et al., Newly produced synaptic vesicle proteins are preferentially used in synaptic transmission. EMBO J, 2018. 37(15).

41. Ratnayaka, A., et al., Extrasynaptic vesicle recycling in mature hippocampal neurons. Nat Commun, 2011. 2: p. 531.

42. Mellman, I., R. Fuchs, and A. Helenius, Acidification of the endocytic and exocytic pathways. Annu Rev Biochem, 1986. 55: p. 663–700.

43. Huotari, J. and A. Helenius, Endosome maturation. EMBO J, 2011. 30(17): p. 3481–500.

44. Haass, C. and D.J. Selkoe, Soluble protein oligomers in neurodegeneration: lessons from the Alzheimer’s amyloid beta-peptide. Nat Rev Mol Cell Biol, 2007. 8(2): p. 101–12.

45. Klein, W.L., G.A. Krafft, and C.E. Finch, Targeting small Abeta oligomers: the solution to an Alzheimer’s disease conundrum? Trends Neurosci, 2001. 24(4): p. 219–24.

46. Soto, C., Unfolding the role of protein misfolding in neurodegenerative diseases. Nat Rev Neurosci, 2003. 4(1): p. 49–60.

47. Taylor, J.P., J. Hardy, and K.H. Fischbeck, Toxic proteins in neurodegenerative disease. Science, 2002. 296(5575): p. 1991–5.

48. Jucker, M. and L.C. Walker, Self-propagation of pathogenic protein aggregates in neurodegenerative diseases. Nature, 2013. 501(7465): p. 45–51.

49. Umeda, T., et al., Intraneuronal amyloid beta oligomers cause cell death via endoplasmic reticulum stress, endosomal/lysosomal leakage, and mitochondrial dysfunction in vivo. J Neurosci Res, 2011. 89(7): p. 1031–42.

50. Mindell, J.A., Lysosomal acidification mechanisms. Annu Rev Physiol, 2012. 74: p. 69–86.

51. Williams, T.L., et al., Abeta42 oligomers, but not fibrils, simultaneously bind to and cause damage to ganglioside-containing lipid membranes. Biochem J, 2011. 439(1): p. 67–77.

52. Butterfield, S.M. and H.A. Lashuel, Amyloidogenic protein-membrane interactions: mechanistic insight from model systems. Angew Chem Int Ed Engl, 2010. 49(33): p. 5628–54.

53. Yerbury, J.J., et al., Walking the tightrope: proteostasis and neurodegenerative disease. J Neurochem, 2016. 137(4): p. 489–505.

54. Ben-Zvi, A., E.A. Miller, and R.I. Morimoto, Collapse of proteostasis represents an early molecular event in Caenorhabditis elegans aging. Proc Natl Acad Sci U S A, 2009. 106(35): p. 14914–9.

55. Jakhria, T., et al., beta2-microglobulin amyloid fibrils are nanoparticles that disrupt lysosomal membrane protein trafficking and inhibit protein degradation by lysosomes. J Biol Chem, 2014. 289(52): p. 35781–94.

56. Bucciantini, M., S. Rigacci, and M. Stefani, Amyloid Aggregation: Role of Biological Membranes and the Aggregate-Membrane System. J Phys Chem Lett, 2014. 5(3): p. 517–27.

57. Lee, S., Y. Sato, and R.A. Nixon, Lysosomal proteolysis inhibition selectively disrupts axonal transport of degradative organelles and causes an Alzheimer’s-like axonal dystrophy. J Neurosci, 2011. 31(21): p. 7817–30.

58. Cataldo, A.M., et al., Endocytic pathway abnormalities precede amyloid beta deposition in sporadic Alzheimer’s disease and Down syndrome: differential effects of APOE genotype and presenilin mutations. Am J Pathol, 2000. 157(1): p. 277–86.

59. Broersen, K., et al., A standardized and biocompatible preparation of aggregate-free amyloid beta peptide for biophysical and biological studies of Alzheimer’s disease. Protein Eng Des Sel, 2011. 24(9): p. 743–50.

60. Williams, T.L., I.J. Day, and L.C. Serpell, The effect of Alzheimer’s Abeta aggregation state on the permeation of biomimetic lipid vesicles. Langmuir, 2010. 26(22): p. 17260–8.

61. Rey, S.A., et al., Ultrastructural and functional fate of recycled vesicles in hippocampal synapses. Nat Commun, 2015. 6: p. 8043.

